# Out-of-the-box bioinformatics capabilities of large language models (LLMs)

**DOI:** 10.1101/2025.08.22.671610

**Authors:** Varsha Rajesh, Geoffrey H. Siwo

## Abstract

Large Language Models (LLMs), AI agents and co-scientists promise to accelerate scientific discovery across fields ranging from chemistry to biology. Bioinformatics- the analysis of DNA, RNA and protein sequences plays a crucial role in biological research and is especially amenable to AI-driven automation given its computational nature. Here, we assess the bioinformatics capabilities of three popular general-purpose LLMs on a set of tasks covering basic analytical questions that include code writing and multi-step reasoning in the domain. Utilizing questions from Rosalind, a bioinformatics educational platform, we compare the performance of the LLMs vs. humans on 104 questions undertaken by 110 to 68,760 individuals globally. GPT-3.5 provided correct answers for 59/104 (58%) questions, while Llama-3-70B and GPT-4o answered 49/104 (47%) correctly. GPT-3.5 was the best performing in most categories, followed by Llama-3-70B and then GPT-4o. 71% of the questions were correctly answered by at least one LLM. The best performing categories included DNA analysis, while the worst performing were sequence alignment/comparative genomics and genome assembly. Overall, LLMs performance mirrored that of humans with lower performance in tasks in which humans had low performance and vice versa. However, LLMs also failed in some instances where most humans were correct and, in a few cases, LLMs excelled where most humans failed. To the best of our knowledge, this presents the first assessment of general purpose LLMs on basic bioinformatics tasks in distinct areas relative to the performance of hundreds to thousands of humans. LLMs provide correct answers to several questions that require use of biological knowledge, reasoning, statistical analysis and computer code.

## Introduction

AI systems have the potential to accelerate every step of the scientific discovery process, including literature search and synthesis, hypothesis generation, research planning, execution of experiments, data collection, analysis and interpretation of results. Recently, there have been numerous studies showcasing the potential of large language models (LLMs), AI agents and co-scientists in accelerating scientific research in fields such as chemistry, material sciences and biology [1-5]. Bioinformatics-computational analysis of DNA, RNA and protein sequencesis central to advancing fundamental biological knowledge and its translation to solutions in health, agriculture and manufacturing. Computational systems that perform bioinformatics analysis using instructions provided in natural language could broaden its impact and accelerate biological research. Bioinformatics analysis is particularly amenable to end-to-end automation with AI systems since it is not constrained by physical, laboratory experimentation.

Several strategies are being pursued in the development of LLMs and AI agents for performing biological research. Most of this work has focused on their capacity for retrieval of information from literature using both general purpose and biological domain specific LLMs trained on scientific literature. For example, BiomedGPT, a multimodal biomedical model, achieved strong benchmark performance using text and image inputs, yet its evaluations focused on classification and QA rather than open-ended reasoning [6]. BioInstruct, another instruction-tuned biomedical LLM, showed improvement on structured QA but did not evaluate performance on realistic, multi-step bioinformatics tasks [7].

A powerful strategy to enable LLMs to perform complex tasks involves providing them access to tools and integrating them using agentic frameworks. Recent reports describe biomedical AI agents and co-scientists capable of reasoning, hypothesis generation, knowledge synthesis and execution of chemistry experiments [1, 4, 8] but only a few have demonstrated capability for automated biological data analysis. For example, TxGemma was obtained by fine-tuning the generalist Gemma model on therapeutic tasks [9] while Google AI co-scientist leverages a multiagent system built on Gemini 2.0 to generate hypotheses [4]. TxAgent performs multi-step reasoning and retrieval of biomedical knowledge across 211 tools with a focus on therapeutic tasks [10], and the Virtual Lab consists of an LLM agent that acts as a principal investigator collaborating with a team of LLM scientist agents with a human in the loop [5]. On the other hand, Biomni, a biomedical AI agent, is capable of mining critical tools, databases and protocols that it subsequently leverages to autonomously perform a wide range of biomedical research tasks [11]. In the case of bioinformatics related tasks, a version of Biomni (A1) with access to 150 specialized biomedical tools, 105 software packages and 59 databases was able to analyze single-cell RNA-seq and ATAQ-seq datasets based on instructions provided in natural language [11]. An alternative approach involves training multi-modal LLMs/AI agents that integrate text and biological sequences. For example, ChatNT is a multi-modal conversational agent for biological sequence analysis tasks, addressing the gap between biological foundation models and conversational agents [12]. BioReason, on the other hand, integrates a DNA foundation model with LLMs to perform multi-step reasoning on specific genomics tasks [13].

While both generalist and biomedical domain LLMs could in principle be integrated with tools for bioinformatics analysis, understanding out-of-the-box capabilities of general purpose LLMs is important for several reasons; i) Generalist LLMs such as OpenAI GPT, Llama, Claude and Gemini models are widely accessible, making them the first go to tools for non-experts; ii) As the world progresses towards artificial general intelligence (AGI), out-of-the-box bioinformatics capabilities of generalist LLMs combined with their improving capacity in a broad range of domains could enable them to discover complex solutions that integrate knowledge within and outside of biomedical sciences-from molecules to ecosystems; iii) Identifying baseline capabilities of general purpose LLMs could also help inform their efficient and reliable integration with biological domain specific tools and agents. Though not a focus of this work, understanding out-of-the box capabilities of general purpose LLMs is critical in anticipating biosecurity risks of these models.

Here, we assess the ability of three generalist LLMs—GPT-3.5, Llama-3-70B, and GPT-4o to solve basic bioinformatics problems and compare their performance to that of humans. We contribute to the growing body of work on AI-assisted bioinformatics by evaluating LLM performance on 104 open-ended questions from Rosalind, an educational platform on practical bioinformatics problems. This study offers insights into both the strengths and limitations of generalist LLMs in performing bioinformatics tasks and advances efforts to make bioinformatics more accessible by reducing the need for specialized expertise.

## Contributions

This work is closely related to recent efforts benchmarking the performance of LLMs in bioinformatics tasks. In particular, the BixBench benchmark dataset consisting of 53 analytical scenarios and 296 open-answer questions curated by bioinformatics experts [14], LabBench which consists of 2400 multi-choice questions on practical biology tasks with a small subset on DNA and protein sequences [15], BaisBench performs assessment of LLMs on single-cell analysis tasks [16] and BioCoder which is a benchmark dataset for evaluating LLMs in bioinformatics-code generation [17]. Key contributions include:

- **Assessment of LLMs vs. thousands of humans on bioinformatics analytical tasks:** We assess performance based on bioinformatics tasks undertaken by thousands of experts and non-experts.
- **Assessment of LLMs on basic, low-level bioinformatics analysis tasks**: We focus on basic/‘low-level’ analytical tasks in a broad range of areas in bioinformatics that include concepts in Mendelian genetics, genetic code translation, genome assembly and phylogenetics.
- **Assessments of LLMs on bioinformatics tasks requiring multi-step reasoning and code writing**: This study is distinct in that it uses bioinformatics analysis problems that require multi-step reasoning, biological context interpretation, and code-like logic to arrive at an exact output.

## Methods Data Sources

This study utilized a total of 104 questions sourced from the Bioinformatics Stronghold section of Rosalind (http://rosalind.info), a widely used online platform that teaches bioinformatics with practice problems. The questions covered key bioinformatics topics of ranging difficulty, including DNA analysis, restriction mapping and enzymes, phylogenetics, combinatorics and probability, RNA and protein translation, genome assembly and graph algorithms, sequence alignment and comparative genomics, genome analysis and computational biology, and motif finding and pattern matching. Each problem included a precise computational task and sample data, some including biological context.

Question difficulty was estimated using the proportion of human users who answered each question correctly, based on Rosalind’s user statistics. These questions had been attempted by between 110 and 68,760 individuals each. The performances of GPT-3.5, Llama 3-70B, and GPT-4o were compared to human performance by evaluating the correctness of their answers against solutions provided by Rosalind. The final output was checked for correctness.

### Large Language Models

We evaluated three large language models: GPT-3.5, GPT-4o (both developed by OpenAI), and Llama-3-70B by Meta. All models were accessed using the University of Michigan’s Generative AI platform (UM-GPT platform), providing a consistent interface. Each question from the Rosalind platform was provided as a prompt to UM-GPT for all three models, keeping the prompt identical between models. Each model was tested on all 104 questions. If a model gave an incomplete response with no final answer, it was prompted once more with the instruction “provide complete answer”. Otherwise, only the first output was recorded.

### Statistical Analysis

All analyses were conducted using Python with the pandas, numpy, matplotlib, seaborn, and scipy libraries. For each of the three models—GPT-3.5, Llama 3 (70B), and GPT-4o. We compared AI-generated answers to the official solutions provided by Rosalind. Model performance was recorded as a binary variable indicating whether the model response was correct (1) or incorrect (0). We categorized each question into overlapping difficulty thresholds based on human accuracy rates: questions where more than 30%, 40%, 50%, 60%, 70%, 80%, and 90% of users answered correctly. Within each benchmark group, we computed and visualized the distribution of AI correctness to assess how model performance varied with question difficulty. To visualize these performance patterns, we created grouped bar charts showing the counts of correct and incorrect AI responses across the benchmark bins. We conducted statistical analyses to evaluate the relationship between human accuracy and model behavior. Additionally, each question was categorized by its primary bioinformatics topic to assess differences in model performance across different content areas.

## Results

### Relative performance of LLMs across all tasks

The Rosalind Bioinformatics Stronghold (http://rosalind.info) consists of 104 questions. We organized these questions into 10 categories consisting of ‘DNA analysis’ (7 questions), ‘Restriction mapping and enzymes’ (2 questions), ‘Phylogenetics’ (14 questions), ‘Combinatorics and probability’ (13 questions), ‘RNA and protein translation’ (13 questions), ‘Genome assembly and graph algorithms’ (9 questions), ‘Sequence alignment and comparative genomics’ (14 questions), ‘Genome analysis/computational biology’ (7 questions), ‘Motif finding and pattern matching’ (12 questions), ‘Genetic inheritance and population genetics’ (10 questions) and a set of 3 questions which we could not assign to a specific topic (‘Unassigned’) (Fig. 1 and Supplementary Table 1). Human performance varied widely, with correctness rates ranging from 25% to 91% for each question. For example, 25% of humans who attempted a question involving ‘Finding All Similar Motifs’ in a sequence were correct, while among the LLMs, only Llama-3-70B was correct. In contrast, 91% of humans provided a correct answer for a question on ‘Counting Disease Carriers’ based on knowledge of the Hardy-Weinberg principle. Both Llama-3-70B and GPT-4o provided correct answers in this question while GPT-3.5 was incorrect. Notably, there was a wide distribution in the number of humans who attempted each question with the most popular question being undertaken by 68,760 humans and least popular by only 110 individuals (median= 1,140; mean = 5,907).

**Fig. 1.**
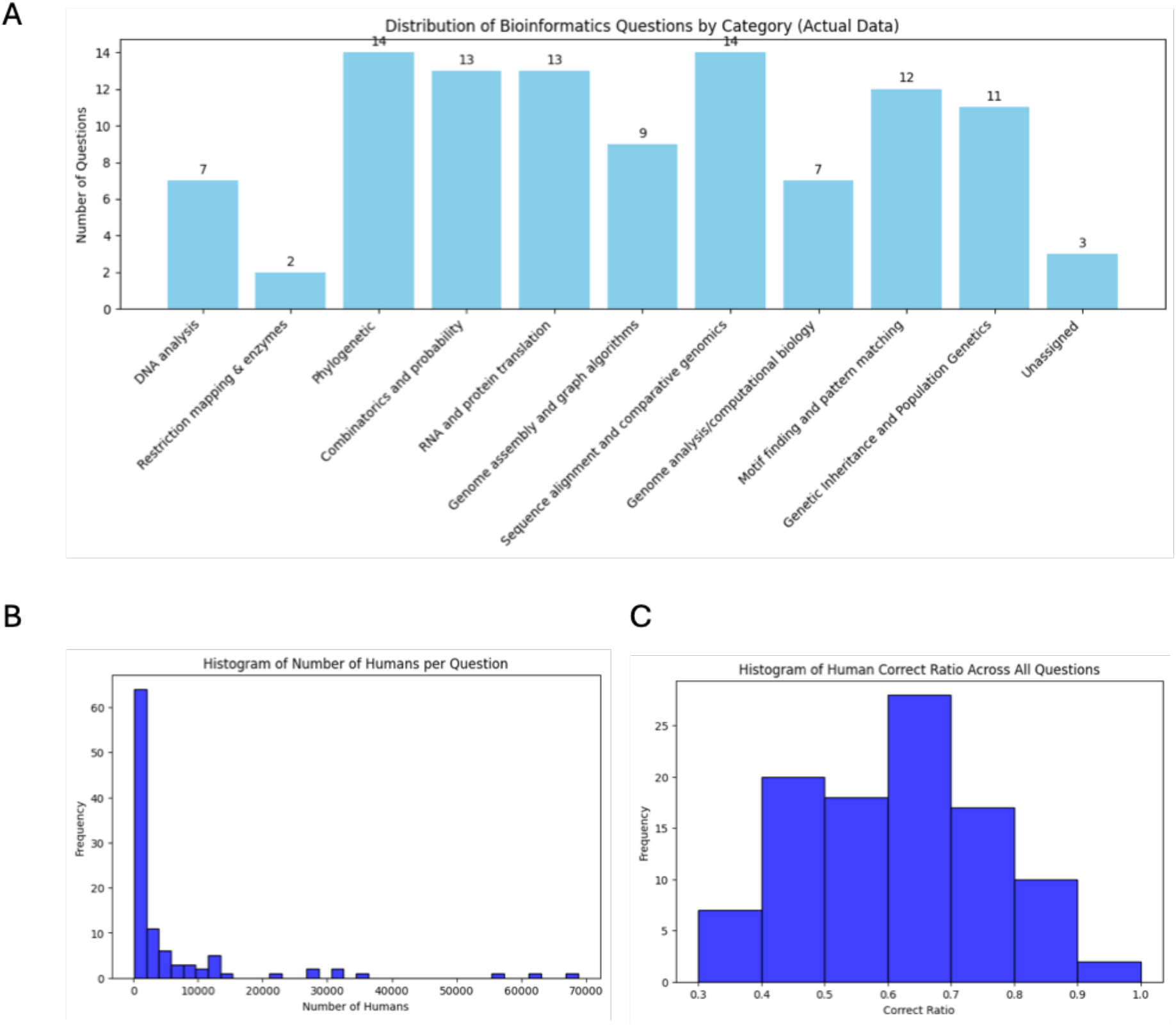
Summary of the Rosalind Bioinformatics Stronghold dataset. (A) Categories of bioinformatics questions represented in the dataset. (B) Number of humans who provided answers per question. (C) Distribution of the average proportion of humans who provided a correct answer to each question (Correct Ratio).

**Fig. 2.**
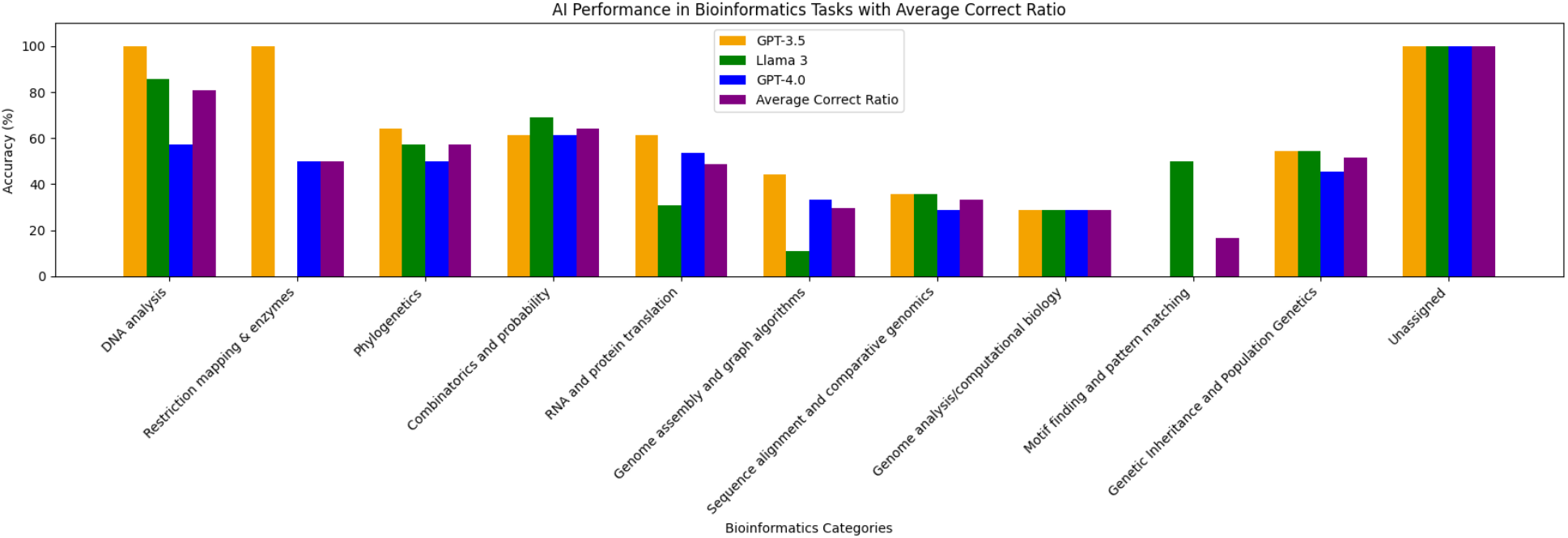
Performance of the LLM models and humans across different categories of bioinformatics tasks. Human performance is based on Average Correct Ratio (average proportion of humans who provided a correct answer for questions in the category).

Overall, GPT-3.5 answered 59/104 of the questions correctly (58%), while Llama-3-70B and GPT-4.0 each answered 49/104 (47%) correctly. 31 questions were answered correctly by all models, and 30 were answered incorrectly by all models (Supplementary Table 1). Thus, in nearly 60% of questions, all the models had the same outcome (i.e. all correct or all incorrect). 71% (74 out of 104 questions) were answered correctly by at least one of the LLMs suggesting that LLMs ensembles or mixture of experts (MoE) could potentially be a strategy for improving performance.

We further analyzed the nature of tasks in which all the LLMs passed or failed. Out of 32 questions in which all LLMs passed, 6 (19%) involved ‘Combinatorics and Probability’, distinct sets of 4 (12%) involved ‘DNA analysis’, ‘Motif finding and pattern matching’, and ‘Phylogenetics’, a separate set of 2 (6%) involved ‘Sequence alignment and comparative genomics’ and Genome analysis/computational biology’. A distinct set of 3 (9%) involved ‘‘Genetic inheritance and population genetics’, ‘Unassigned’ and ‘RNA and protein translation’.

Out of 31 questions in which all LLMs failed, separate sets of 3 (10%) involved ‘Combinatorics and Probability’, ‘Genetic inheritance and population genetics’, ‘Genome analysis/computational biology’, a distinct set of 5 (16%) involved ‘Genome assembly and graph algorithms’ and ‘Motif finding and pattern matching’. A distinct set of 2 (6%) involved ‘Phylogenetics’, ‘RNA and protein translation’, and 7 (23%) involved ‘Sequence alignment and comparative genomics’.

We investigated whether the performance of humans in tasks in which LLMs provide correct answers differs from that in tasks in which LLMs provide incorrect answers. Human performance tended to be higher in tasks in which LLMs provided correct answers compared to in tasks where LLMs were incorrect (Table 1).

**Table 1.** LLM and human performance.

|  | *GPT-3.5* | *Llama 3* | *GPT-4o* | *All LLMs* |
| --- | --- | --- | --- | --- |
| ***Average proportion of correct humans in AI correct*** | 0.64 | 0.62 | 0.63 | 0.62 |
| ***Average proportion of correct humans in AI incorrect*** | 0.59 | 0.61 | 0.60 | 0.59 |

To assess relationships between models in terms of their correctness, we determined cases where pairs of models showed similar patterns of correctness where one model was incorrect. GPT-3.5 and Llama-3-70B had the most overlap (11 overlapping correct answers in cases where GPT-4o was incorrect), followed by GPT-3.5 and GPT-4o (8 overlapping correct answers), and finally GPT-4o and Llama had the least overlap (3 overlapping correct answers). Overlapping correct answers for GPT-3.5 and Llama-3-70B were in ‘Phylogenetics’ (3 questions answered correctly only by the two models), ‘Sequence alignment and comparative genomics’ (1 overlap), ‘Combinatorics and probability’ (1 overlap), ‘RNA and protein translation’ (1 overlap), ‘Genetic inheritance and population genetics’ (2 overlaps), ‘DNA analysis’ (2 overlaps) and ‘Genome analysis/Computational biology’ (1 overlap). In the case of GPT-3.5 and GPT-4o, questions where only these 2 models gave the correct answer involved ‘Sequence alignment and comparative genomics’ (2 overlaps), ‘Genome assembly and graph algorithms’ (2 overlaps), ‘Restriction enzymes and mapping’ (1 overlap), ‘RNA and protein translation’ (1 overlap), ‘Combinatorics and probability’ (1 overlap) and ‘Genetic inheritance and population genetics’ (1 overlap). GPT-4o and Llama-3-70B had overlapping correct answers in ‘Combinatorics and probability’ (1 overlap), ‘Phylogenetics’ (1 overlap) and ‘Genetic inheritance and population genetics’ (1 overlap).

Next, we asked whether certain questions were only correctly answered by one model. There were 10 questions in which only GPT-3.5 provided correct answers compared to 6 for GPT-4o and 5 for Llama. For the questions where GPT-3.5 was the only model with correct answers, 3 involved ‘RNA and protein translation’, 2 involved ‘Phylogenetics’, 1 question each involved ‘Restriction mapping and enzymes’, ‘Genome assembly and graph algorithms’, ‘Genome analysis/computational biology’, ‘DNA analysis’, ‘Motif finding and pattern matching’. GPT-4o only correct answers involved 2 questions in ‘Phylogenetics’ and 1 question each in ‘Motif finding and pattern matching’, Genome analysis/computational biology’, ‘Genetic inheritance and population genetics’, and ‘RNA and protein translation’. Llama-3-70B only correct answers involved 2 questions in ‘Sequence alignment and comparative genomics’ and 1 question each from ‘Combinatorics and probability’, Genome analysis/computational biology’, and ‘Motif finding and pattern matching’.

Top-performing categories across the 3 models were ‘DNA analysis’ (GPT-3.5: 100%, Llama 3: 85.7%, GPT-4o: 57.1%), ‘Unassigned’ (100% for all models), and ‘Restriction mapping and enzymes’ (GPT-3.5: 100%). Notably, there were very few questions in these categories which biases the results. Lower-performing areas included ‘Genome assembly and graph algorithms’ (GPT-3.5: 44.4%), ‘Sequence alignment’ (GPT-3.5 & Llama 3: 35.7%), and ‘Motif finding and pattern matching’ (GPT-3.5 & GPT-4o: 0%).

### Performance of LLMs at different benchmarks of human difficulty

We next investigated the performance of LLMs in tasks in which humans had high vs. low performance. The top 20 questions in which humans had the highest performance (average of 82% of humans correct) involved ‘Genetic inheritance and population genetics’ (4 questions), ‘DNA analysis’ (2 questions), ‘Combinatorics and probability’ (3 questions), ‘Motif finding and pattern matching’ (1 question), ‘RNA and protein analysis’ (1 question), ‘Restriction mapping and enzymes’ (1 question), ‘Phylogenetics’ (1 question), ‘Unassigned’ (1 question). Among these top 20 tasks in humans, all the three LLMs were correct in 5 cases (25%) and incorrect in 4 cases (20%). GPT-3.5 was correct in 11 cases (55%), GPT-4o in 10 cases (50%), and Llama-3-70B in 8 cases (40%). The tasks in which all LLMs were incorrect yet humans had high performance involved questions asking about inferring genotype from pedigree (86% out of 363 humans correct), modeling random genomes (83% out of 4994 humans correct), the founder effect and genetic drift (80% out of 375) and computing a maximum alignment score when aligning two given DNA sequences (80% of 178).

In contrast, the top 20 questions in which humans had the lowest performance (average 41% humans provided correct answers) involved: ‘Motif finding and pattern matching’ (1 question), ‘Sequence alignment and comparative genomics’ (7 questions), ‘Phylogenetics’ (4 questions), ‘RNA and protein translation’ (2 questions), ‘Genome analysis/Computational biology’ (1 question), and ‘Combinatorics and probability’ (2 questions). Among these tasks where humans had low performance, all the three LLMs were correct in 3 cases and incorrect in 8 cases. GPT-3.5 was correct in 7 cases, GPT-4o in 4 cases, and Llama-3-70B in 7 cases. Within the cases where all the LLMs were correct were a question on determining the global alignment between two input protein sequences given a constant gap penalty (38% out of 406 humans correct), determining total number of non-crossing perfect matchings of base pair edges in an RNA secondary structure (43% out of 1,542 humans correct) and, finding a consensus DNA sequence and profile matrix for a set of provided DNA sequences (46% out of 15,250 humans correct).

We hypothesized that the number of humans who undertook a given question is another dimension that reflects the potential difficulty of the questions to humans. On average, 64% of humans provided correct answers among the top 20 most attempted questions compared to 52% correct answers in the 20 least attempted questions. This implies that human tendency to undertake the questions likely reflects perceived level of difficulty. Therefore, we next assessed the performance of LLMs based on this perceived level of difficulty.

Among the top 20 most attempted questions by humans, all the three LLMs provided the correct answers in 11 cases. All LLMs were incorrect in only 2 cases. Notably, the two questions in which all LLMs failed in this set involved ‘Finding a Motif in DNA’ (attempted by 28,720 humans with 72% providing the correct answer) and translation of ‘Open Reading Frames’ (attempted by 7,780 humans with 45% getting it correct). GPT-3.5 provided 16 correct answers in the top 20 most attempted questions, GPT-4o had 13 correct answers, and Llama had 14 correct answers. In contrast, for the 20 least attempted questions by humans, all the LLMs were incorrect on 11 cases and correct on only 1 case. The single correct case was on ‘Linguistics Complexity of a Genome’, a question attempted by 264 individuals with 73% of them providing correct answers. GPT-3.5 provided 6 correct answers in the set of least attempted questions, GPT-4o provided 3 correct answers, and Llama provided 6 correct answers. This suggests that LLMs have a lower ability to solve questions that would be considered difficult by non-expert bioinformaticians. LLMs may find such questions challenging potentially due to the lack of sufficient examples in their training datasets.

To better understand the relationship between LLM and human performance, we established different benchmarks of human performance based on the proportion of humans who provided a correct response to each question. GPT-3.5 had more correct than incorrect responses at benchmarks from 30% to 79%, while GPT-4o and Llama-3-70B provided more incorrect answers at most benchmarks except for the 70 to 79% benchmark (Fig. 3).

**Fig. 3.**
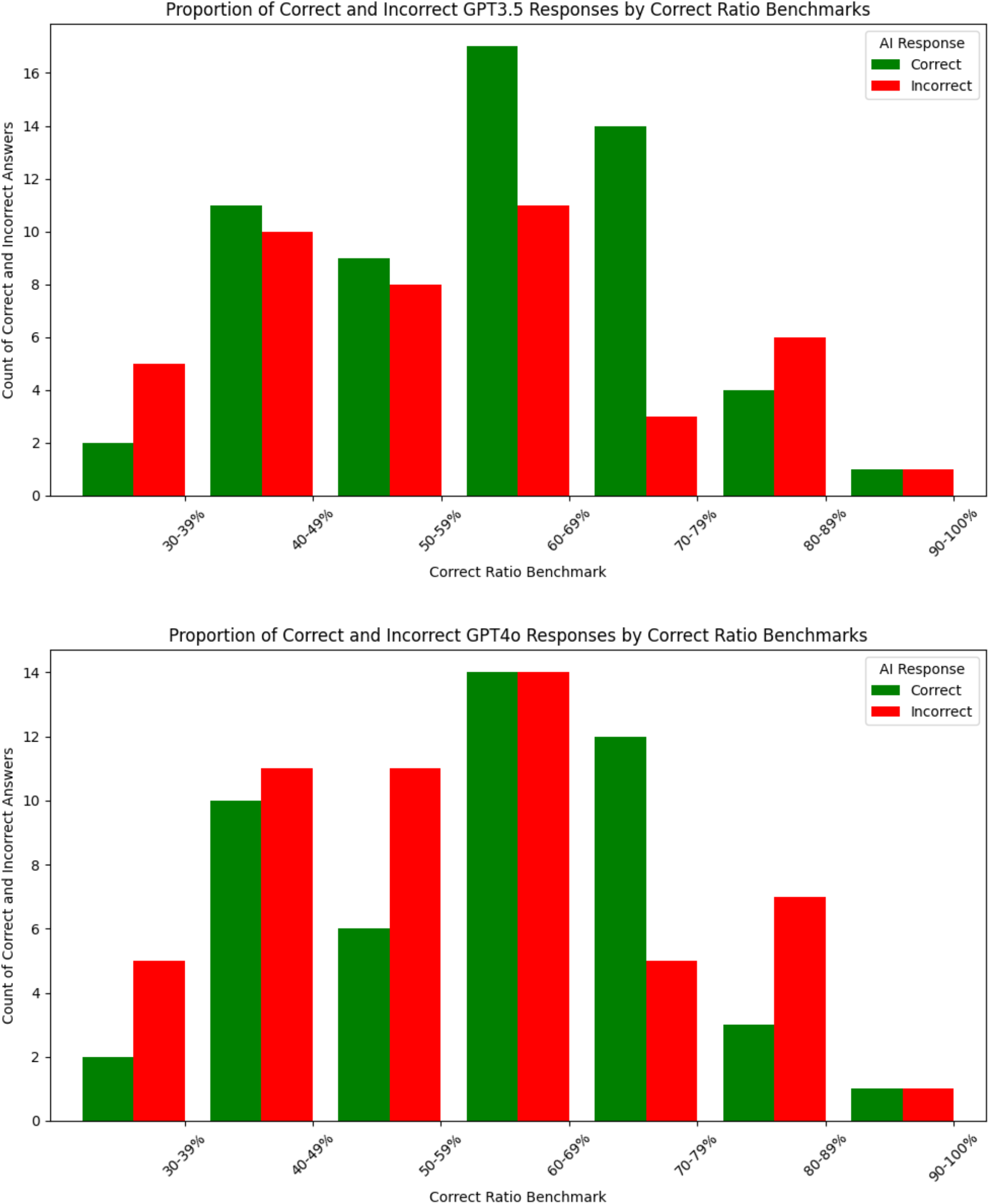
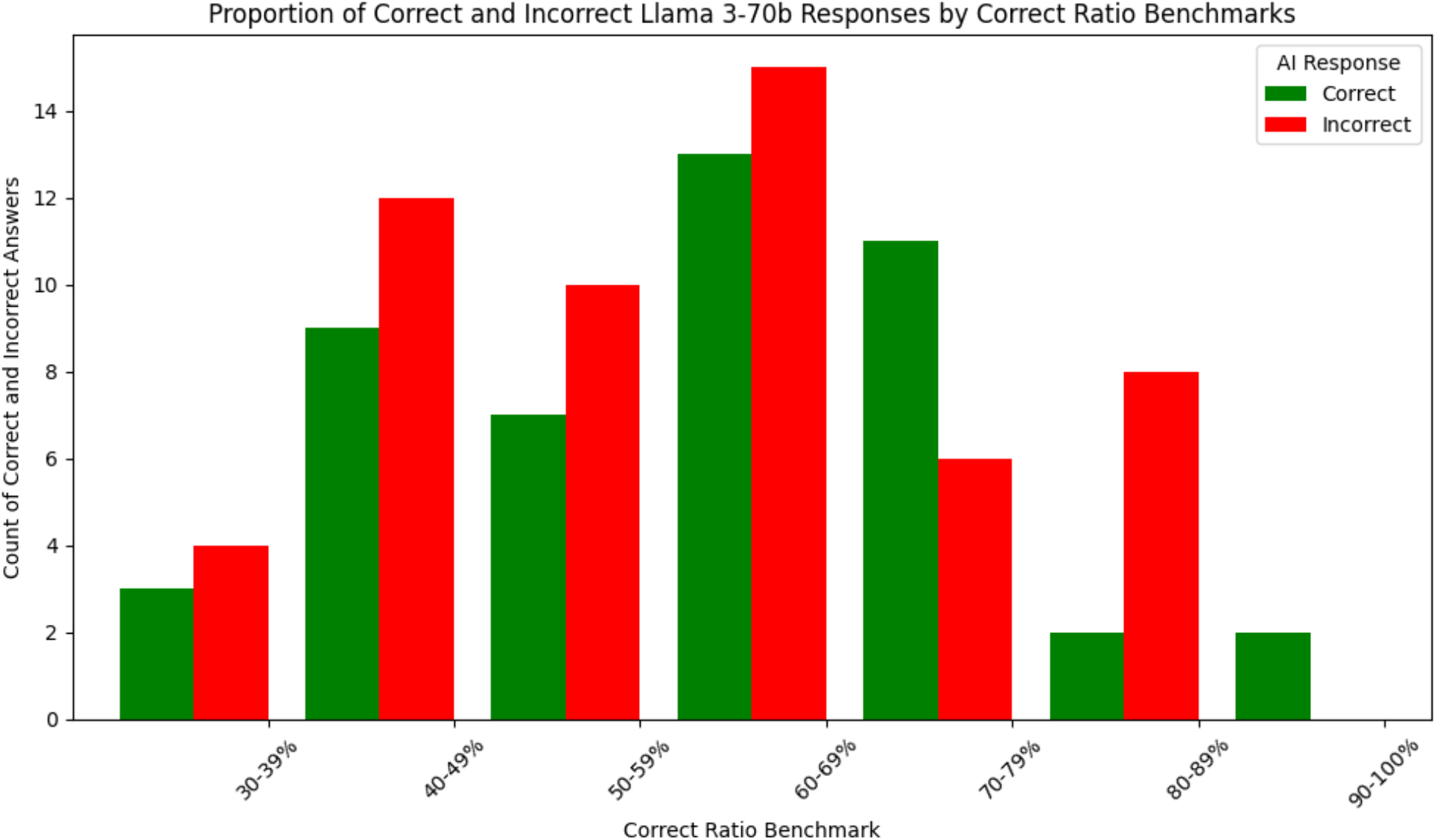
Performance of LLMs at different benchmarks of human performance.

## Discussion and Conclusion

Understanding the bioinformatics analysis capacity of general purpose LLMs is an important endeavor given the broad accessibility of these models to biomedical researchers and the generical public. Furthermore, given the natural language capabilities of general purpose LLMs, out-of-the-box capabilities in these models could enhance their integration with specialized biomedical AI agents and bioinformatics tools while benefiting from rapid progress in non-biomedical LLMs. Despite the explosion of biomedical domain LLMs and AI agents, most of these systems are developed and evaluated for specialized biomedical tasks, do not take natural language inputs, and are intended for domain experts, limiting their broad utility.

This study explored the capabilities of the large language models GPT-3.5, Llama 3-70B, and GPT-4.0 in performing bioinformatics tasks relative to humans with a wide range of expertise levels. While each model correctly answered 49 to 58% of the 104 open-ended questions sourced from the Rosalind Bioinformatics Stronghold, the models also failed in some questions where most humans obtained the correct answer. It is notable that the performance of LLMs was very dependent on the performance of humans on specific tasks-tasks in which humans had high performance were well done by the LLMs compared to those in which humans had low performance. A dependency on the number of humans who undertook a given question was also evident with LLMs showing high performance on tasks that were undertaken by many humans compared to performance in tasks that fewer humans undertook. Thus, it appears that the number of humans who undertook a given question reflects the perceived difficulty of the question.

LLMs demonstrated strong performance in areas such as DNA analysis and restriction mapping— domains that often involve more straightforward computations or well-defined rules. Conversely, they struggled in categories that require higher-order reasoning, algorithm selection, or pattern recognition, such as genome assembly, phylogenetics, motif finding, and sequence alignment. Despite these limitations, the models exhibited notable proficiency in generating answers that required a combination of biological reasoning, statistical analysis, and computational logic. Their ability to solve complex problems without prior specialization in bioinformatics underscores the potential for LLMs to assist in this domain, particularly for educational or exploratory use cases. However, the models’ occasional production of incomplete or incorrect answers highlights current gaps in training and the need for improved domain adaptation.

This study is limited by the fact that the bioinformatics tasks presented are not comprehensive enough to determine whether one LLM is better than the other. While GPT-3.5 had the highest performance, this may not be generalizable to performance on other bioinformatics tasks. It is important to note that the tasks in Rosalind require multi-step reasoning using different aspects of biological knowledge. Thus, although the number of questions tested is relatively small, the extent of bioinformatics knowledge covered by the questions is much larger. Future work will focus on extending the evaluations to include the latest versions of LLMs and more bioinformatics tasks, assessing reasoning traces/chain-of-thought, enhancing LLM performance using mixture of experts, and investigating biosecurity risks.

## Data and Code Availability

All code used for data preprocessing, analysis, and visualization was written in Google Colab. The complete analysis was implemented in Python. https://github.com/SiwoResearch/lllm-bioinformatics.

Supplementary Table 1 contains results of comparisons of the three models across 104 questions

